# Oxygen-sensing neurons reciprocally regulate peripheral lipid metabolism via neuropeptide signaling in *Caenorhabditis elegans*

**DOI:** 10.1101/142422

**Authors:** Rosalind Hussey, Emily Witham, Erik Vanstrum, Jaleh Mesgarzadeh, Harkaranveer Ratanpal, Supriya Srinivasan

**Affiliations:** Department of Molecular Medicine, Department of Neuroscience and Dorris Neuroscience Center, The Scripps Research Institute, 10550 North Torrey Pines Road, La Jolla, CA 92037, USA; Current Address: Janssen Pharmaceutical Companies of Johnson and Johnson, 3210 Merryfield Row, San Diego, CA 92121, USA; Department of Biology, University of California, San Diego, 9500 Gilman Drive, La Jolla, CA 92093, USA; Current Address: Liberty University College of Osteopathic Medicine, 306 Liberty View Lane, Lynchburg, VA 24502, USA

## Abstract

The mechanisms by which the sensory environment instructs metabolic homeostasis remains poorly understood. In this report, we show that oxygen, a potent environmental signal, is an important regulator of whole body lipid metabolism. *C. elegans* oxygen-sensing neurons reciprocally regulate peripheral lipid metabolism under normoxia in the following way: under high oxygen and food absence, URX sensory neurons are activated, and stimulate fat loss in the intestine, the major metabolic organ for *C. elegans*. Under lower oxygen conditions or when food is present, the BAG sensory neurons respond by repressing the resting properties of the URX neurons. A genetic screen to identify modulators of this effect led to the identification of a BAG-neuron-specific neuropeptide called FLP-17, whose cognate receptor EGL-6 functions in URX neurons. Thus, BAG sensory neurons counterbalance the metabolic effect of tonically active URX neurons via neuropeptide communication. The combined regulatory actions of these neurons serve to precisely tune the rate and extent of fat loss, to the availability of food and oxygen.

## Introduction

The central nervous system plays a critical role in regulating whole body energy balance. In mammals, in addition to the role of the hypothalamus, there is good evidence showing that the sensory nervous system also regulates whole body metabolism (1-4). However, the identification of discrete sensory modalities and the underlying signaling pathways that regulate metabolism in peripheral tissues has remained a tremendous challenge. As a result, basic questions regarding the molecular and physiological mechanisms underlying the neuronal control of lipid metabolism remain unknown even as rates of obesity, diabetes and their complications soar worldwide.

In the nematode *Caenorhabditis elegans*, many ancient mechanisms of sensory and metabolic regulation have been preserved, thus offering an opportunity to dissect the pathways by which the nervous system regulates lipid metabolism. The *C. elegans* nervous system is relatively well-defined both genetically and anatomically (5, 6), thus genes underlying discrete sensory modalities can be rapidly assessed for roles in whole body metabolism. The intestine is the seat of metabolic control, occupies the majority of the animal, and is the predominant depot for fat storage and mobilization. Thus, changes in whole body metabolism can be effectively encapsulated by monitoring metabolic readouts in the intestine. *C. elegans* sensory neurons play pivotal roles in regulating metabolism (7, 8). At least 3 discrete sensory inputs: food availability (9-11), population density (12) and environmental oxygen (13) are relayed from sensory neurons to the intestine. These sensory inputs instruct the duration and magnitude of fat loss, effectively coupling environmental information with body fat metabolism via neuroendocrine hormones. The molecular and neuroendocrine mechanisms by which sensory information is relayed from the nervous system to the intestine has begun to shed light on the interaction between genetic mechanisms and environmental conditions in controlling lipid metabolism.

Neuronal oxygen sensing is a sensory modality that is richly informative to *C. elegans*, which grow in environments replete with bacteria, their major food source. Respiring bacteria drop the local ambient concentration of oxygen from 21% (atmospheric) to a range between 10-13%. Worms presented with an oxygen gradient from 0-21% choose a range between 10-13%, and avoid higher and lower oxygen concentrations, the better to remain in areas that reflect the balance between sufficient food and oxygen (14, 15). Importantly, these responses are distinct from hypoxia-related responses, which occur at 3% oxygen and below, are encoded by the conserved hypoxia genes including HIF-1, and are distinct from mechanisms of oxygen sensing during normoxia (16). Oxygen preference within the normoxic range is encoded by two pairs of sensory neurons (17). The URX neurons detect high (21%) oxygen via a guanylate cyclase called GCY-36, and initiate an aversive behavioral response (17, 18). On the other hand, BAG neurons detect low (5-10%) oxygen via an inhibitory guanylate cyclase called GCY-33, and also initiate aversion (17, 19). Thus, URX and BAG neurons function together to retain animals in food environments that contain an optimal balance of food and oxygen.

In a screen for neuronal regulators of fat metabolism, we uncovered a role for the URX neurons in regulating oxygen-dependent fat loss (13). When fasted, body fat stores are metabolized for the production of energy. A time course of food withdrawal at atmospheric (21%) oxygen shows that the intestinal fat stores of *C. elegans* are steadily depleted, and by 3 hours young adults lose ~80% of their body fat stores; near-complete fat loss in the intestine occurs by 4 hours (Fig. 2 *A*; 13). Wild-type animals exposed to 10% oxygen retain approximately twice as much body fat as those exposed to high oxygen. We previously showed that this effect is not regulated by changes in food intake or locomotor rates at high versus low oxygen (13). Rather, it occurs because of a selective metabolic shift towards fat utilization that is differentially regulated by oxygen availability. In the intestine, this metabolic shift depends upon the conserved triglyceride lipase called ATGL-1, which is transcriptionally regulated by oxygen. Thus, animals exposed to high oxygen show greater fat loss via ATGL-1 activation, than those exposed to low oxygen for the same duration. This differential metabolic response to oxygen depends on neuronal oxygen sensing via the URX neurons, and functions via the oxygen - sensing guanylate cyclase GCY-36. We found that a gustducin-like Gα protein called GPA-8 functions as a negative regulator of GCY-36, and serves to limit the tonic, constitutive activity of URX neurons by repressing resting Ca^2+^ levels. Loss of GPA-8 disinhibits the tonic activity of the URX neurons at 10% oxygen (when they are normally silent), which leads to increased fat loss (13).

**Fig. 1.**
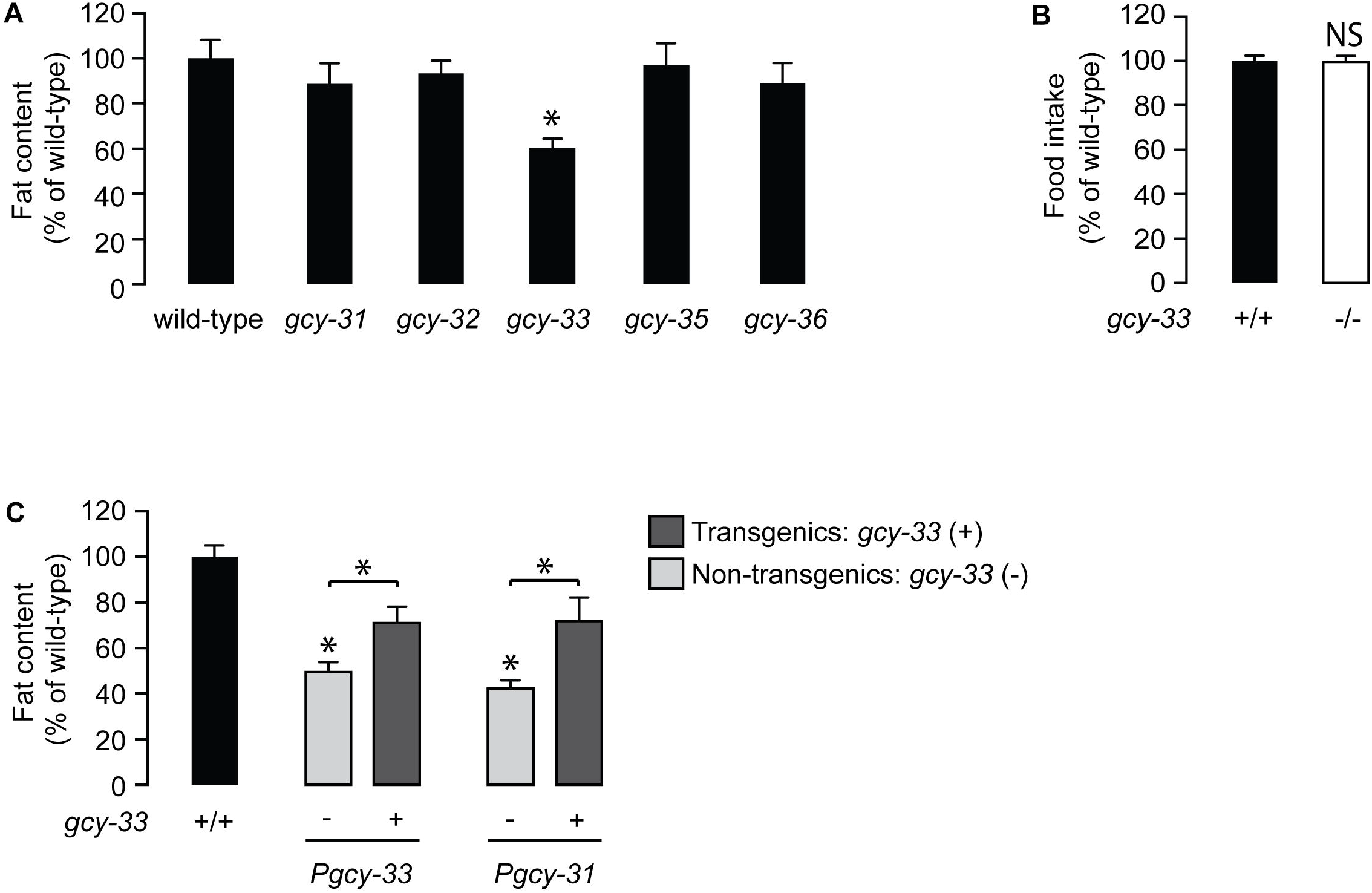
The oxygen sensor GCY-33 acts in BAG neurons to regulate lipid metabolism in intestinal cells. (*A*) Animals were fixed and stained with Oil Red O. Fat content was quantified for each genotype and is expressed as a percentage of wild-type animals ± SEM (n=12-20). *, p<0.05 by one-way ANOVA. See also Fig. S1. (B) Food intake in *gcy-33* mutants expressed as a percentage of wild-type ± SEM (n=10). NS, not significant by Student’s t-test. (C) For each transgenic line bearing *gcy-33* expression using the indicated promoter, non-transgenic animals (−) and transgenic animals (+) are shown. Relative to non-transgenic controls (light gray bars), transgenic animals (dark gray bars) bearing the *gcy-33* and heterologous *gcy-31* promoters restore body fat content to that seen in wild-type animals. Data are expressed as a percentage of body fat in wild-type animals ± SEM (n=18-20). *, p<0.05 by one-way ANOVA.

**Fig. 2.**
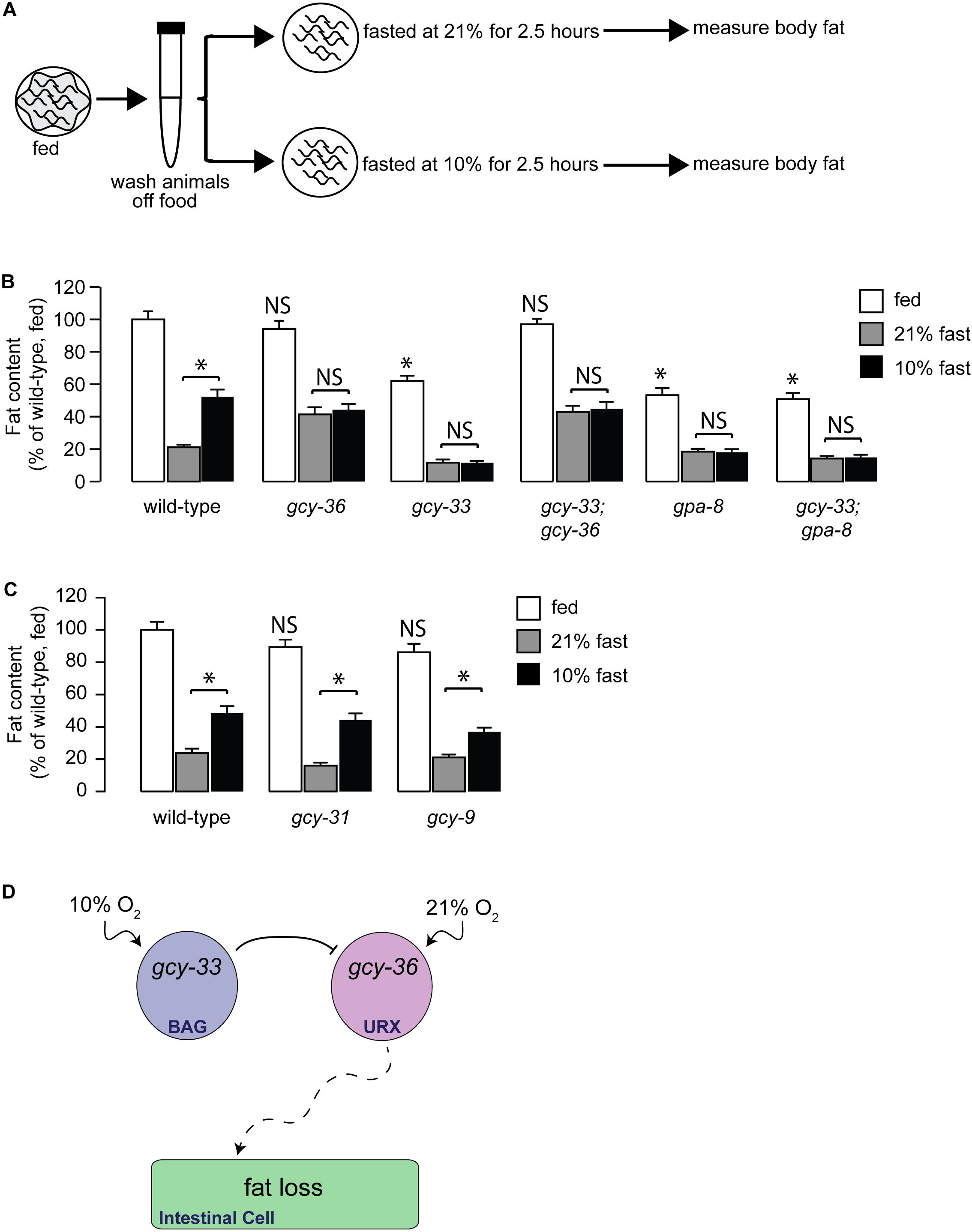
GCY-33 acts upstream of the URX-cGMP signaling pathway in fat regulation. (*A*) Schematic depiction of the oxygen-dependent fat loss assay, reprinted with permission from Elsevier (13). Worms were washed off food, and then fasted. Fasted worms were exposed to either 21% (high) oxygen or 10% (low) oxygen. At the end of the fasting period, worms were fixed and stained with Oil Red O. (*B*) Worms of the indicated genotypes were subjected to the oxygen-dependent fat loss assay described in (*A*). Fat content was quantified for each genotype and condition. Data are expressed as a percentage of body fat in wild-type fed controls ± SEM (n=16-20). NS, not significant and *, p<0.05 by one-way ANOVA. (*C*) Worms of the indicated genotypes were subjected to the oxygen-dependent fat loss assay. Fat content was quantified for each genotype and condition. Data are expressed as a percentage of body fat in wild-type fed controls ± SEM (n=18-20). NS, not significant and *, p<0.05 by one-way ANOVA. (*D*) Schematic depiction of the neuroendocrine axis through which sensation of low oxygen concentrations by the guanylate cyclase GCY-33 in the BAG neurons inhibits URX-mediated utilization of fat in the intestinal cells.

Our findings on the role of the URX neurons in regulating oxygen-dependent fat loss made us contemplate a potential role for the BAG neurons, which reciprocally regulate behaviors associated with sensing low oxygen (5-10%) and more broadly, of the role of normoxic oxygen in regulating lipid metabolism (17, 19). The URX and BAG neurons are activated at high and low oxygen, respectively (14, 15, 17, 19-21). However, these neurons are known to be tonic sensors of oxygen, suggesting that they would need to be kept in the ‘off state’ by active repression. Neuronal mechanisms underlying repression of tonically active neurons has not been described. In this study, we report that BAG neurons play an important role in regulating oxygen-dependent fat loss. We find that BAG neurons relay information about low oxygen to the URX neurons and inhibit them to maintain low tonic activity. The physiological consequence of this neuronal signaling is to reduce the rate and extent of fat loss in the intestine, during low oxygen. In this context, communication from BAG neurons to URX neurons occurs via the FLP-17 neuropeptide from the BAG neurons and its cognate receptor, the G protein coupled receptor EGL-6, which functions in the URX neurons. Thus, we find that BAG neurons play a critical role in regulating the rate and extent of fat loss via modulation of the resting properties of URX neurons. Our work points to a model in which neurons with opposing sensory roles establish a mutually reinforcing circuit via neuropeptide modulation, and that the neuronal sensation of low oxygen serves as a negative regulator of fat loss.

## Results

### The soluble guanylate cyclase GCY-33 regulates intestinal lipid metabolism from the oxygen-sensing BAG neurons

Because of our previously-described role for the URX-neuron-specific soluble guanylate cyclase GCY-36 in regulating oxygen-dependent fat loss (13), we examined all available null mutants of the soluble guanylate cyclase (sGC) family for changes in body fat (Fig. 1 *A* and Fig. S1 *A* and *B*). Relative to wild-type animals, *gcy-33* mutants showed a ~40% reduction in body fat stores, whereas loss of other sGC mutants did not show an appreciable difference. Food intake (Fig. 1 *B*) and locomotor rates (17) are indistinguishable between wild-type and *gcy-33* mutants, suggesting that differences in the metabolism of fat stores underlies the fat phenotype of *gcy-33* mutants. *gcy-33* mutants are defective in the behavioral and neuronal responses to low oxygen, which are encoded by the BAG neurons (17, 19). In addition to the BAG neurons, GCY-33 is also reported to be expressed in the URX neurons (17, 22). To evaluate its site of action in regulating body fat, we restored *gcy-33* cDNA in the *gcy-33* mutants under its own promoter, and using heterologous promoters for the BAG and URX neurons. We observed robust partial rescue of body fat stores upon re-expression of GCY-33 under its endogenous promoter and in the BAG neurons (Fig. 1 *C*), but did not observe meaningful rescue of body fat stores with its reexpression in the URX neurons (data not shown). Incomplete rescue of *gcy-33* mutants with exogenous cDNA has been previously observed, for reasons that are currently not known (17). Regardless, our experiments indicate that GCY-33 functions in the BAG neurons to regulate body fat stores.

### GCY-33 signaling from BAG neurons regulates oxygen-dependent fat loss via URX neuron signaling

We measured the extent to which *gcy-33* mutants played a role in oxygen-dependent fat loss. Fasted wild-type adults initiate fat loss in an oxygen-dependent manner: wild-type animals exposed to 21% (henceforth, ‘high’) oxygen metabolize approximately twice as much body fat as those exposed to 10% (henceforth, ‘low’) oxygen (Fig. 2 *A* and *B*). We previously showed that this effect is not regulated by changes in food intake or locomotor rates at high versus low oxygen (13). Rather, it occurs because of a selective metabolic shift towards fat utilization that is differentially regulated by oxygen availability. Thus, animals exposed to high oxygen show greater fat loss than those exposed to low oxygen for the same duration. As expected, *gcy-36* mutants show complete suppression of fat loss at high oxygen, consistent with the previously-observed role for the URX neurons in responses to high oxygen (Fig. 2 *B*). We found that *gcy-33* mutants showed the opposite phenotype to the *gcy-36* mutants: they had greater fat loss at low oxygen than wild-type animals, and were indistinguishable from those exposed to high oxygen. This result is consistent with loss of BAG neuron activity in *gcy-33* mutants (17), and indicates a role for BAG neurons in suppressing fat loss at low oxygen (Fig. 2 *B*). The observed effects of the BAG neurons on oxygen-dependent fat loss were specific to *gcy-33* signaling, because null mutants of two other BAG-specific guanylate cyclases *gcy-31* and *gcy-9* (23) did not suppress oxygen-dependent fat loss (Fig. 2 *C*).

We were surprised to note that removal of the URX-specific sGC *gcy-36* fully suppressed the phenotype of *gcy-33* single mutants, such that the *gcy-33;gcy-36* mutants resembled the *gcy-36* single mutants alone (Fig. 2 *B*). Because gcy-36 functions in the URX neurons and not in the BAG neurons (17), this result suggested that the effect of GCY-33 signaling from the BAG neurons was dependent on GCY-36 signaling from the URX neurons (Fig. 2 *D*). The increased fat loss of *gcy-33* mutants under low oxygen (Fig. 2 *B*) was highly reminiscent of the *gpa-8* mutants which we had previously studied (Fig. 2 *B*; 13). GPA-8 is a gustducin-like Gα protein which functions within the URX neurons themselves, and serves to inhibit GCY-36. *gpa-8* mutants have reduced fat stores in the fed state, show increased fat loss at low oxygen, and are indistinguishable from those fasted at high oxygen. *gcy-33;gpa-8* double mutants resembled each single mutant alone, with no additive effects (Fig. 2 *B*). We previously showed that the fat-regulatory property of *gpa-8* mutants arises from increased constitutive activation of the URX neurons because of *gcy-36* de-repression (13), again resembling the *gcy-33;gcy-36* mutants. Together, the data suggests that *gcy-33* signaling from BAG neurons negatively regulates URX neurons and oxygen-dependent fat loss (Fig. 2 *D*).

### GCY-33 modulates the resting state of URX neurons

To directly study the effects of GCY-33 signaling on URX function, we measured neuronal activity using Ca^2+^ imaging in living animals. We used the genetically-encoded calcium indicator GCaMP5k as a reporter for URX activity which has been optimized for greater sensitivity to threshold activation properties (13, 24). Wild-type animals bearing the GCaMP5k transgene expressed under the URX-specific promoter *flp-8* showed robust calcium influx at 21% oxygen (Fig. 3 *A* and *C*), as previously described (13, 25). We observed two properties of URX activation in *gcy-33* mutants crossed into the *flp-8∷GCaMP5k* transgenic line. First, at 21% oxygen there was an approximately 30% decrease in maximal activation of URX neurons in *gcy-33* mutants compared to wild-type animals (Fig. 3 *B* and *D* and *E*). Second, at 10% oxygen, we observed an approximately two-fold increase in baseline (F_0_) fluorescence values in *gcy-33* mutants compared to wild-type animals (Fig. 3 *F*). Importantly, we verified that there was no observed difference in *flp-8* promoter activity at 10% oxygen (Fig. 3 *G*). When measuring URX peak responses to the oxygen upshift in absolute GCaMP5k fluorescence levels, no significant difference between wild-type animals and *gcy-33* mutants was observed (Fig. 3 *H*). The major effect of GCY-33 therefore lies in controlling Ca^2+^ concentrations in the URX neurons at 10% oxygen. These data were highly reminiscent of the effect of *gpa-8* mutants on URX Ca^2+^ dynamics (13). Together, our data indicate that BAG neurons, via *gcy-33* signaling, inhibit resting URX properties at low (10%) oxygen (Fig. 2 *D*).

**Fig. 3.**
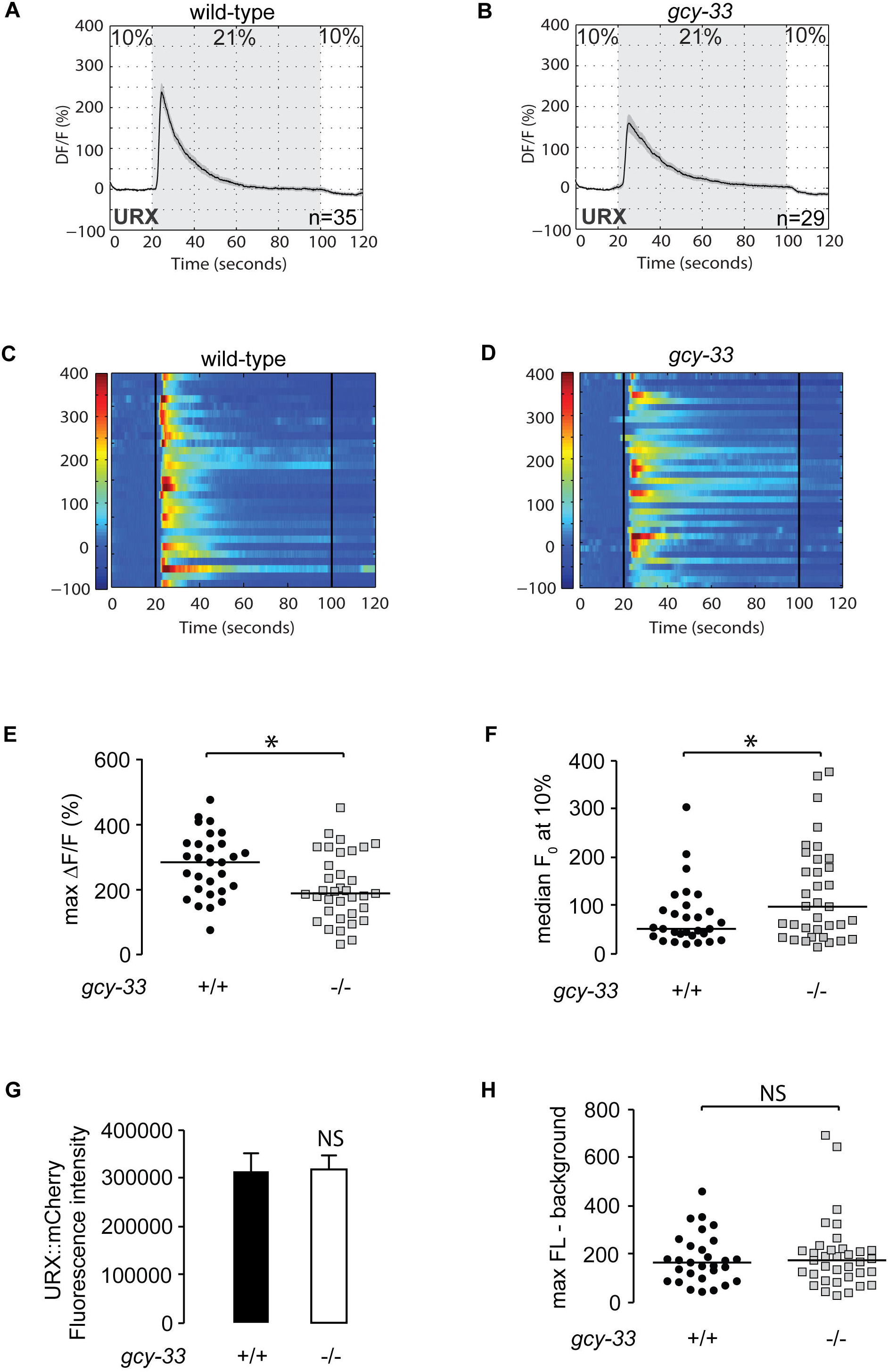
GCY-33 controls the resting-state Ca^2+^ concentrations in URX neurons. (*A-D*) Measurements of neuronal activity by Ca^2+^ imaging of URX neurons for each genotype. The number of animals used for each condition is shown in the figure. We conducted Ca^2+^ imaging experiments in URX neurons in living wild-type animals and *gcy-33* mutants bearing *GCaMP5k* under the control of the *flp-8* promoter. Oxygen concentrations in the microfluidic chamber were 10% (low) and 21% (high) as indicated. (*A-B*) For each genotype, black traces show the average percent change of GCaMP5k fluorescence (ΔF/F_0_) and gray shading indicates SEM. (*C-D*) Individual URX responses are shown for each genotype; each row represents one animal. (*E*) Maximum ΔF/F_0_ values are shown for individual wild-type animals and *gcy-33* mutants. Bars indicate the average value within each genotype. *, p<0.05 by Student’s t-test. (*F*) Individual baseline fluorescence (F_0_) values at 10% (low) oxygen are shown for individual wild-type animals and *gcy-33* mutants. Bars indicate the median value within each genotype. *, p<0.05 by Student’s t-test. (*G*) We imaged mCherry fluorescence in wild-type animals and *gcy-33* mutants expressing both GCaMP5k and mCherry under the control of the *flp-8* promoter. Images were taken in animals exposed to 10% (low) oxygen. For each genotype, the fluorescence intensity was imaged at the same exposure, determined to be within the linear range. Fluorescence intensity was quantified and expressed as an average ± SEM (n=20). NS, not significant by Student’s t-test. (*H*) The background-subtracted maximum fluorescence (max FL) at 21% (high) oxygen is shown for each wild-type animal and *gcy-33* mutant. Bars indicate the median value within each genotype. NS, not significant by Student’s t-test.

### Neuropeptide communication from BAG neurons to URX neurons coordinates oxygen-dependent fat loss

BAG neurons are not known to form synaptic connections with the URX neurons. In addition, null mutants of the dense core vesicle-specific activator protein CAPS/UNC-31 that regulate neuropeptide secretion (26-28) blocked oxygen-dependent fat loss, whereas null mutants of the synaptic vesicle protein UNC-13 that regulates conventional neurotransmitter release (29, 30), did not (Fig. S2). To explore a potential neuropeptide-based mechanism of communication from the BAG neurons to the URX neurons, we conducted a targeted genetic screen of all available mutants of the neuropeptide gene families (*flp, nlp* and *ins* gene families, 77/113 genes). Our top hit was a neuropeptide called FLP-17, the canonical BAG-neuron-specific neuropeptide (31-33) (Fig. 4 *A*). Interestingly, *flp-17* null mutants were nearly identical to the *gcy-33* mutants in oxygen-dependent fat loss. That is, relative to wild-type animals, fed *flp-17* mutants had reduced fat stores, showed increased fasting-dependent fat loss at low oxygen, and were indistinguishable from those at high oxygen. In addition, *gcy-33;flp-17* and *gpa-8;flp-17* double mutants resembled either single mutant alone (Fig. 4 *A*), suggesting that GCY-33 and FLP-17 from BAG neurons function in a linear pathway with GPA-8 signaling in URX neurons to inhibit fat loss at low oxygen.

**Fig. 4.**
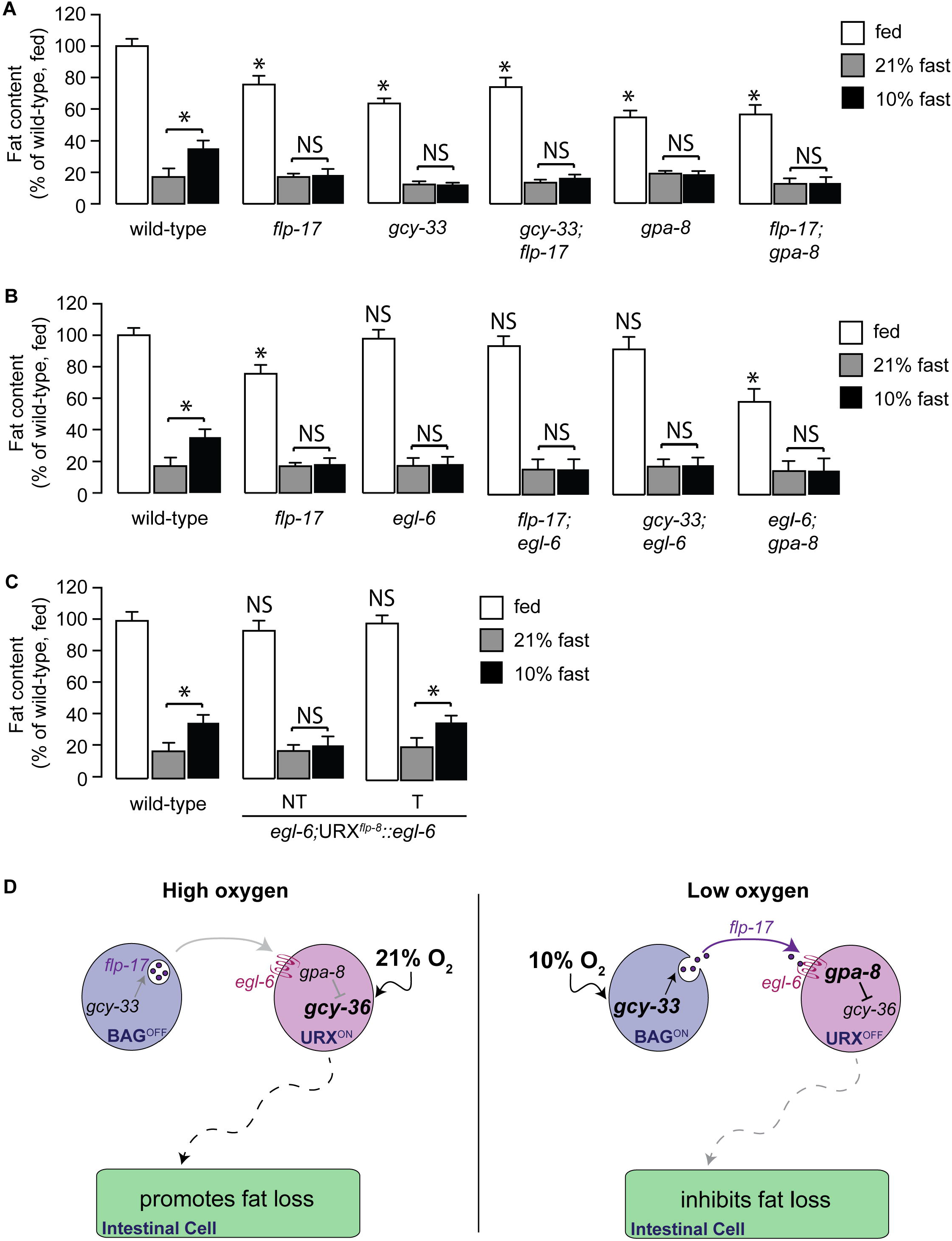
Neuropeptidergic signaling drives fat mobilization through oxygen sensing. (*A,B*) Worms of the indicated genotypes were subjected to the oxygen-dependent fat loss assay. Fat content was quantified for each genotype and condition. Data are expressed as a percentage of body fat in wild-type fed controls ± SEM (n=16-20). NS, not significant and *, p<0.05 by one-way ANOVA. (*C*) Worms from transgenic line bearing *egl-6* expression using the *flp-8* promoter were subjected to the oxygen-dependent fat loss assay. Non-transgenic animals (−) and transgenic animals (+) are shown. Relative to non-transgenic controls (light gray bars), transgenic animals (dark gray bars) restore body fat content at 10% (low) oxygen to that seen in wild-type animals. Data are expressed as a percentage of body fat in wild-type animals ± SEM (n=16-20). NS, not significant and *, p<0.05 by one-way ANOVA. (*D*) Model depicting the role of the oxygen-sensing neurons in intestinal lipid metabolism. When animals are exposed to high (21%) oxygen (left panel), the URX neurons are activated via the soluble guanylate cyclase GCY-36, which promotes fat loss in the intestine (13). Under low (10%) oxygen (right panel), signaling via GCY-33 in the BAG neurons promotes the release of the neuropeptide FLP-17, which is detected by EGL-6 in the URX neurons. Communication from BAG neurons to the URX neurons, mediated by FLP-17/EGL-6 signaling dampens URX neuron activity, thus limiting fat loss. Our model provides a mechanism by which tonic activity of the URX neurons is controlled, and the symbiotic relationship between the BAG neurons and the URX neurons that converts the perception of oxygen to tune the rate and extent of fat loss in the intestine.

A GPCR called EGL-6 functions as the cognate receptor for the FLP-17 neuropeptide in the control of egg-laying (33). Activation of the EGL-6 receptor inhibits egg-laying via its function in the HSN neurons, whereas loss-of-function mutants do not have an egg-laying phenotype. We found that null mutants of the *egl-6* gene did not have overt defects in body fat stores, but were defective in oxygen-dependent fat loss indistinguishably from the *flp-17* mutants: loss of *egl-6* led to increased fat loss at low oxygen such that they resembled fat loss at high oxygen (Fig. 4 *B*). *flp-17;egl-6* double mutants resembled the *egl-6* single mutants in all respects: in the fed state they did not show an appreciable difference in body fat stores, and showed increased fat loss in low oxygen, without additive effects. This result suggested that FLP-17 and EGL-6 function in a linear genetic pathway to regulate oxygen-dependent fat loss.

*egl-6* expression had been noted in *C. elegans* head neurons (33). Because our signaling pathway indicated a role of communication from the BAG to the URX neurons via the FLP-17 neuropeptide, we tested whether EGL-6 functions in the URX neurons. In *egl-6* null mutants, we restored expression in the URX neurons and measured oxygen-dependent fat loss. Relative to non-transgenic animals, *egl-6* re-expression in URX neurons fully restored oxygen-dependent fat loss (Fig. 4 *C*), suggesting that EGL-6 functions in the URX neurons to limit fat loss at low oxygen. *gcy-33;egl-6* and *egl-6;gpa-8* double mutants did not yield additive effects relative to each single mutant alone, suggesting that these genes function in a linear pathway to regulate oxygen-dependent fat loss (Fig. 4 *D*).

## Discussion

In this report, we show that the *C. elegans* oxygen-sensing neurons reciprocally regulate peripheral lipid metabolism. Under lower oxygen conditions, the BAG sensory neurons respond by repressing the URX neurons, which are tonic sensors of higher oxygen. Molecularly, repression is effected by a neuropeptide and its cognate receptor. Thus, BAG sensory neurons counterbalance the metabolic effect of the tonically active URX neurons. The physiological consequence of this repression is to limit fat utilization when oxygen levels are low.

In previous work we had defined the role of the URX neurons in stimulating fat loss in response to high environmental oxygen (Fig. 4 *D*, left panel). When oxygen levels are high, and food supplies dwindle, URX neurons are activated by the actions of GCY-36, ultimately leading to increased fat loss via activation of the ATGL-1 lipase (13). In the present study we define one mechanism by which the URX neurons are held in the ‘off state’ under conditions of low oxygen, that is, via repression from BAG neurons. It has been appreciated for some years that URX neurons are tonic sensors of environmental oxygen, suggesting that they must be turned off by an active repression, the molecular genetics of which remained unknown. Our model suggests that BAG neurons in *C. elegans* play an essential role in mediating tonic repression of URX-neuron resting properties in the following way: the sensation of low oxygen concentrations by the guanylate cyclase GCY-33 in the BAG neurons initiates the release of the neuropeptide, FLP-17, which is detected via the GPCR EGL-6 in the URX neurons. FLP-17/EGL-6 signaling leads to decreased basal activity of URX neurons via the GPA-8-mediated repression of GCY-36. Thus, in low oxygen, URX neurons are retained in the ‘off state’, ultimately limiting fat loss (Fig. 4 *D*, right panel).

Collectively, URX neurons are held in the off state in two ways. First, by internal sensing of body fat stores (13) and second, by the presence of food or low oxygen via the BAG neurons (this study). We suggest that it is a combination of decreased GCY-33 activation in BAG neurons and increased GCY-36 activation in URX neurons that increases fat utilization in the intestinal cells. Our experiments reveal an important relationship between the BAG and URX neurons in the regulation of metabolic homeostasis. In doing so, we have identified a novel pair of fat regulating neurons, and a physiological context in which the counterbalancing actions of BAG and URX neurons play a vital role.

## Materials and Methods

### Animal Maintenance and Strains

*C. elegans* was cultured as described (34). N2 Bristol, obtained from the Caenorhabditis Genetic Center (CGC) was used as the wild-type reference strain. The mutant and transgenic strains used are listed in Table S1. Animals were synchronized for experiments by hypochlorite treatment, after which hatched L1 larvae were seeded on plates with the appropriate bacteria. All experiments were performed on day 1 adults.

### Cloning and Transgenic Strain Construction

Promoters and genes were cloned using standard PCR techniques from N2 Bristol worm lysates or cDNA and cloned using Gateway Technology™ (Life Technologies). Promoter lengths were determined based on functional rescue and are available upon request. All rescue plasmids were generated using polycistronic GFP. Transgenic rescue strains were constructed by microinjection into the *C. elegans* germline followed by visual selection of transgenic animals under fluorescence. For the microinjections, 5-10 ng/μl of the desired plasmid was injected with 25 ng/μl of an *unc-122::GFP* coinjection marker and 65-70 ng/μl of an empty vector to maintain a total injection mix concentration of 100 ng/μl. In each case, 10–20 stable transgenic lines were generated. Two lines were selected for experimentation based on consistency of expression and transmission rate. For GCaMP5k transgenic animals, 5 ng/μl of *Pflp-8::GCaMP5k* was injected with 2 ng/μl of a *Pflp-8::mCherry* coinjection marker.

### Oil Red O Staining

Oil Red O staining was performed as described (13). Within a single experiment, roughly 3,500 animals were fixed and stained, 100 animals were visually inspected on slides, following which 15-20 animals were imaged for each genotype/condition. All experiments were repeated at least 3 times.

### Image Acquisition and Quantitation

Black and white images of Oil Red O stained animals and fluorescent images were captured using a 10X objective on a Zeiss Axio Imager microscope. Lipid droplet staining in the first four pairs of intestinal cells was quantified as described (9). Within each experiment, approximately 14-20 animals from each condition were quantified. Images were quantified using ImageJ software (NIH).

### Food Intake

Food intake was measured by counting pharyngeal pumping, as previously described (35). For each animal, the rhythmic contractions of the pharyngeal bulb were counted over a 10 s period under a Zeiss M2 Bio Discovery microscope. For each genotype, 10 animals were counted per condition and the experiment was repeated at least three times.

### Oxygen-dependent Fat Loss Assay

The experiments were conducted as described (13). For each strain, approximately 3,500 synchronized L1 larvae were seeded onto each of three plates. Worms were grown at 20°C for 48 h after which all plates were transferred to the bench top. Worms subjected to the fasting protocol were washed off the plates with PBS in 5 sequential washes over a 30-minute period to eliminate residual bacteria, and then seeded onto NGM plates without food. Worms were then subjected to a 2.5 h fasting period at either 21% or 10% oxygen. To establish the time course of fasting, pilot experiments were conducted at atmospheric (21%) oxygen. The “21% fasted” plates were placed in a non-airtight container at room temperature. The “fed” control plates were placed in a similar but separate container. The “10% fasted” plates were placed in a custom-designed sealed acrylic oxygen chamber (TSRI Instrumentation and Design Lab), fitted with inlet and outlet valves. The inlet valve was connected via bubble tubing to a pressurized oxygen and nitrogen pre-mixture containing 10% oxygen (Praxair, Inc.), and the outlet valve was exposed to air. All plates were positioned right side up without lids. The sealed chamber was then perfused for 15 min with 10% oxygen. Following perfusion, both valves were closed. During the experiment, pressure inside the chamber was held constant, as judged by a gauge placed inside the oxygen chamber. The chamber was kept at room temperature for an additional 2.25 h, so that all fasted conditions remained off food for a total of 2.5 h following the washes. At the end of this period, worms from the respective conditions were collected for Oil Red O staining.

### Calcium Imaging

We used a microfluidic chamber constructed with the oxygen-permeable poly(dimethylsiloxane) (PDMS) as described (17). A Valvebank II (AutoMate Scientific, Inc.) was used to control input from two pressurized pre-mixtures of oxygen and nitrogen containing either 10% oxygen or 21% oxygen (Praxair, Inc.). The gas flow rate was set to 0.26 psi at the outlet of the chamber as judged by a VWR^™^ traceable pressure meter. Immediately before imaging, individual day 1 adult animals were sequentially transferred to two unseeded plates. Individual *C. elegans* adults were then transported into the chamber in a drop of S Basal buffer containing 6mM levamisole (Acros Organics B.V.B.A.) via Tygon tubing (Norton). Animals were constantly submerged in S Basal buffer while inside the chamber. After the animals were immobilized inside the chamber, GCaMP5k fluorescence was visualized at 40x magnification using a spinning disk confocal microscope (Olympus) using MetaMorph™ (version 6.3r7, Molecular Devices). Worms were pre-exposed to 10% oxygen for 5 min in the microfluidic chamber as described (17). GCaMP5k fluorescence was recorded by stream acquisition for two minutes at a rate of 8.34 frames/second, with an exposure time of 20 ms using a 12-bit Hamamatsu ORCA-ER digital camera. Each animal was recorded once. GCaMP5k-expressing neurons were marked by a region of interest (ROI). The position of the ROI was tracked using the “Track Objects” function in MetaMorph™. An adjacent ROI was used to subtract background from the total integrated fluorescence intensity of the ROI. Data were analyzed using MATLAB (MathWorks, Inc.). Fluorescence intensity is presented as the percent change in fluorescence relative to the baseline (ΔF/F_0_). F_0_ was measured in worms exposed to 10% oxygen during the first 9-13 seconds for each recording and calculated as an average over that period. All animals were day1 adults at the time of imaging. The number of animals used for each condition is denoted in the figures.

### Statistics

Wild-type animals were included as controls for every experiment. Error bars represent SEM. Student’s t-test, one-way ANOVA, and two-way ANOVA were used as indicated in the figure legends.

## Acknowledgments

This work was supported by research grants to SS from the NIH/NIDDK (R01 DK095804), Novartis Advanced Discovery Institute and the Ray Thomas Edwards Foundation. We are grateful to the Knockout Consortium at Tokyo Women’s Medical University. Some strains were provided by the CGC, which is funded by the NIH Office of Research Infrastructure Programs (P40 OD010440). We thank Dr. Manuel Zimmer (IMP, Vienna) for critical advice and reagents for calcium imaging, and Dr. Kathryn Spencer, Dorris Neuroscience Center, The Scripps Research Institute for assistance with Ca^2+^ imaging. We also thank Dr. Nicole Littlejohn and other members of the Srinivasan lab for critical comments on the manuscript.

## Author Contributions

RH, EW and SS designed the experiments. RH and SS wrote the manuscript. RH and EW conducted the experiments, with assistance from HR, EV and JM. All authors read and approved the manuscript.

## Supporting Information

**Fig. S1. *gcy-33* null mutants exhibit a significant decrease in body fat content.** (*A*) Images of wild-type animals and *gcy-33* mutants fixed and stained with Oil Red O. Animals are oriented facing upwards with the pharynx at the top of each image. For each genotype, images depict the full range of the observed phenotype. (*B*) The integrated density of the lipid droplets is used to quantify body fat stores, as described in the Materials and Methods. Graph represents the integrated density values of individual wild-type animals and *gcy-33* mutants. *, p<0.05 by Student’s t-test.

**Fig. S2. Neuropeptidergic signaling drives fat mobilization through oxygen sensing.** Worms of the indicated genotypes were subjected to the oxygen-dependent fat loss assay. Fat content was quantified for each genotype and condition. Data are expressed as a percentage of body fat in wild-type fed controls ± SEM (n=20). NS, not significant and *, p<0.05 by one-way ANOVA.

**Table S1. *C. elegans* strains used in this study.**

## References

1. Mykytyn K, et al. (2002) Identification of the gene (BBS1) most commonly involved in Bardet-Biedl syndrome, a complex human obesity syndrome. Nature genetics 31(4):435–438.

2. Cao L, et al. (2011) White to brown fat phenotypic switch induced by genetic and environmental activation of a hypothalamic-adipocyte axis. Cell metabolism 14(3):324–338.

3. Linford NJ, Kuo T-H, Chan TP, & Pletcher SD (2011) Sensory Perception and Aging in Model Systems: From the Outside In. Annual Review of Cell and Developmental Biology 27(1):759–785.

4. Srinivasan S (2015) Regulation of Body Fat in Caenorhabditis elegans. Annual Review of Physiology 77(1).

5. Larsch J, Ventimiglia D, Bargmann CI, & Albrecht DR (2013) High-throughput imaging of neuronal activity in Caenorhabditis elegans. Proceedings of the National Academy of Sciences 110(45):E4266–E4273.

6. Zaslaver A, et al. (2015) Hierarchical sparse coding in the sensory system of Caenorhabditis elegans. Proceedings of the National Academy of Sciences of the United States of America 112(4):1185–1189.

7. Greer ER, Pérez CL, Van Gilst MR, Lee BH, & Ashrafi K (2008) Neural and Molecular Dissection of a C. elegans Sensory Circuit that Regulates Fat and Feeding. Cell Metabolism 8(2):118–131.

8. Neal SJ, et al. (2016) A Forward Genetic Screen for Molecules Involved in Pheromone-Induced Dauer Formation in Caenorhabditis elegans. G3 (Bethesda, Md.) 6(5):1475–1487.

9. Noble T, Stieglitz J, & Srinivasan S (2013) An Integrated Serotonin and Octopamine Neuronal Circuit Directs the Release of an Endocrine Signal to Control C. elegans Body Fat. Cell Metabolism 18(5):672–684.

10. Palamiuc L, et al. (2017) A tachykinin-like neuroendocrine signalling axis couples central serotonin action and nutrient sensing with peripheral lipid metabolism. Nature Communications 8:14237.

11. Srinivasan J, et al. (2008) A blend of small molecules regulates both mating and development in Caenorhabditis elegans. Nature 454(7208):1115–1118.

12. Hussey R, et al. (2017) Pheromone-sensing neurons regulate peripheral lipid metabolism in Caenorhabditis elegans. PLoS genetics 13(5):e1006806.

13. Witham E, et al. (2016) C. elegans body cavity neurons are homeostatic sensors that integrate fluctuations in oxygen availability and internal nutrient reserves. Cell reports 14(7):1641–1654.

14. Cheung BH, Cohen M, Rogers C, Albayram O, & de Bono M (2005) Experience-dependent modulation of C. elegans behavior by ambient oxygen. Current biology: CB 15(10):905–917.

15. Chang AJ, Chronis N, Karow DS, Marletta MA, & Bargmann CI (2006) A distributed chemosensory circuit for oxygen preference in C. elegans. PLoS biology 4(9):e274.

16. Jiang H, Guo R, & Powell-Coffman JA (2001) The Caenorhabditis elegans hif-1 gene encodes a bHLH-PAS protein that is required for adaptation to hypoxia. Proceedings of the National Academy of Sciences 98(14):7916–7921.

17. Zimmer M, et al. (2009) Neurons detect increases and decreases in oxygen levels using distinct guanylate cyclases. Neuron 61(6):865–879.

18. Gray JM, et al. (2004) Oxygen sensation and social feeding mediated by a C. elegans guanylate cyclase homologue. Nature 430(6997):317–322.

19. Skora S & Zimmer M (2013) Life(span) in balance: oxygen fuels a sophisticated neural network for lifespan homeostasis in C. elegans. Embo j 32(11):1499–1501.

20. Cheung BH, Arellano-Carbajal F, Rybicki I, & de Bono M (2004) Soluble guanylate cyclases act in neurons exposed to the body fluid to promote C. elegans aggregation behavior. Current Biology 14(12):1105–1111.

21. Busch KE, et al. (2012) Tonic signaling from O2 sensors sets neural circuit activity and behavioral state. Nature neuroscience 15(4):581–591.

22. Yu S, Avery L, Baude E, & Garbers DL (1997) Guanylyl cyclase expression in specific sensory neurons: a new family of chemosensory receptors. Proceedings of the National Academy of Sciences 94(7):3384–3387.

23. Brandt JP & Ringstad N (2015) Toll-like receptor signaling promotes development and function of sensory neurons required for a C. elegans pathogen-avoidance behavior. Current Biology 25(17):2228–2237.

24. Akerboom J, et al. (2012) Optimization of a GCaMP Calcium Indicator for Neural Activity Imaging. The Journal of Neuroscience 32(40):13819.

25. Schrödel T, Prevedel R, Aumayr K, Zimmer M, & Vaziri A (2013) Brain-wide 3D imaging of neuronal activity in Caenorhabditis elegans with sculpted light. Nat Meth 10(10):1013–1020.

26. Walent JH, Porter BW, & Martin TFJ (1992) A novel 145 kd brain cytosolic protein reconstittesu Ca2+-regulated secretion in permeable neuroendocrine cells. Cell 70(5):765–775.

27. Berwin B, Floor E, & Martin TFJ (1998) CAPS (Mammalian UNC-31) Protein Localizes to Membranes Involved in Dense-Core Vesicle Exocytosis. Neuron 21(1):137–145.

28. Lin X-G, et al. (2010) UNC-31/CAPS docks and primes dense core vesicles in C. elegans neurons. Biochemical and biophysical research communications 397(3):526–531.

29. Richmond JE, Davis WS, & Jorgensen EM (1999) UNC-13 is required for synaptic vesicle fusion in C. elegans. Nature neuroscience 2(11):959–964.

30. Madison JM, Nurrish S, & Kaplan JM (2005) UNC-13 Interaction with Syntaxin Is Required for Synaptic Transmission. Current Biology 15(24):2236–2242.

31. Li C, Nelson LS, Kim K, Nathoo A, & Hart AC (1999) Neuropeptide gene families in the nematode Caenorhabditis elegans. Annals of the New York Academy of Sciences 897(1):239–252.

32. Kim K & Li C (2004) Expression and regulation of an FMRFamide-related neuropeptide gene family in Caenorhabditis elegans. Journal of Comparative Neurology 475(4):540–550.

33. Ringstad N & Horvitz HR (2008) FMRFamide neuropeptides and acetylcholine synergistically inhibit egg-laying by C. elegans. Nature neuroscience 11(10):1168–1176.

34. Brenner S (1974) The genetics of Caenorhabditis elegans. Genetics 77(1):71–94.

35. Srinivasan S, et al. (2008) Serotonin regulates C. elegans fat and feeding through independent molecular mechanisms. Cell metabolism 7(6):533–544.

